# Septins Provide Microenvironment Sensing and Cortical Actomyosin Partitioning in Motile Amoeboid T Lymphocytes

**DOI:** 10.1101/2022.01.18.476840

**Authors:** Alexander S. Zhovmer, Alexis Manning, Chynna Smith, Pablo J. Sáez, Xuefei Ma, Denis Tsygankov, Alexander X. Cartagena-Rivera, Rakesh K. Singh, Erdem D. Tabdanov

## Abstract

The all-terrain motility of lymphocytes in tissues and tissue-like gels is best described as amoeboid motility. For amoeboid motility, lymphocytes do not require specific biochemical or structural modifications to the surrounding extracellular matrix. Instead, they rely on changing shape and steric interactions with the microenvironment. However, the exact mechanism of amoeboid motility remains elusive. Here we report that septins shape T cells for amoeboid motility. Specifically, septins form F-actin and alpha-actinin-rich cortical rings at the sites of cortex-indenting collisions of T cells with the extracellular matrix. Cortical rings compartmentalize cells into chains of spherical segments that are spatially conformed to the available lumens, forming transient ‘hourglass’-shaped steric locks onto the surrounding collagen fibers. The steric lock facilitates pressure-driven peristaltic propulsion of cytosolic content by individually contracting cell segments. Our results demonstrate that septins provide microenvironment-guided partitioning of actomyosin contractility and steric pivots for amoeboid motility of T cells in tissue-like microenvironments.

**GLOSSARY:**

1. Steric interactions - interactions by the means of their spatial collision dependent on objects’ shapes.
2. Steric guidance - cell navigation within crowded 3D environments, determined by the available passages around and between steric hindrances.
3. Peristaltic treadmilling - locomotion mode by the means of a repeated sequence of polarized cell cortex extension, stabilization, and retraction, accompanied by translocation of nucleus and cytoplasm *via* circumferential cortex contractility.

**Significance Statement:** T cells can be highly motile, searching for cognate antigens or better yet targets in chimeric antigen receptor therapy settings. However, mechanisms of motility remain elusive for T cells migrating in structurally and biochemically diverse tissues. Here we address one pivotal question of basic and clinical immunology - How T cells achieve the ‘all-terrain’ motility? Here we decipher and report septin-based T cell motility in a 3D tissue-like environment. Specifically, we show that septins facilitate cell morphological responsiveness to the steric obstacles, *i.e.*, collagen fiber-wise partitioning of actomyosin cortex contractility and cell-obstacle steric interactions. These responses coordinate peristaltic propulsion of the lymphocyte’s cytosolic content along its individually contracting cell segments, forming the obstacle-avoiding motility, *i.e.,* circumnavigation, shared across various tested lymphocytes.

## INTRODUCTION

Amoeboid motility is the most typical mode of lymphocyte locomotion in tissues. However, its exact mechanisms are not completely understood. For example, T cells do not require specific biochemical or structural modifications to the surrounding collagen matrix for amoeboid motility. Instead, they rely on the active changing of cell shape and steric (*i.e.,* non-adhesive) interactions with a collagen matrix, which allows them to circumnavigate, *i.e*., steer around the collagen fibers, while finding available lumens for passage (Sadjadi et al., 2020; Wolf et al., 2003). Moreover, it is also not clear why the observed efficiency of amoeboid motility is proportional to the contractility of cortical actomyosin and inversely proportional to the cell-microenvironment adhesive interactions across all sterically interactive environments, such as nano-topographic surfaces, 3D collagen matrices, and cancer tumor tissue samples (Krummel et al., 2014; Tabdanov et al., 2021; Takesono et al., 2010).

The current model of amoeboid motility suggests that actomyosin contractility stimulates the formation of cell blebs, which serve as precursors for cell extension (Charras and Paluch, 2008). During blebbing, the actomyosin cortex and plasma membrane serve both as a substrate for bleb formation and as a primary generator of cytoplasmic hydrostatic pressure, which causes repeating events of destabilization and transient ruptures of binding interactions between the underlying cortex and plasma membrane (Charras et al., 2006, 2005; Fritzsche et al., 2013). Conversely, the blebbing model of amoeboid motility does not include alternative mechanisms of non-adhesive locomotion described for immune cells, *e.g.,* cell sliding by the retrograde cortex treadmilling in the ‘leading bleb’ (Adams et al., 2018; Renkawitz et al., 2009), and the motility by the active cortical deformation (Reversat et al., 2020). The blebbing model also does not address the spatiotemporally coordinated 3D extension of the cell cortex within a structurally discrete microenvironment, which is required for effective directional propulsion of immune cells.

Therefore, considering clinical interest, there are significant scientific efforts aimed at improving our understanding of amoeboid lymphocyte locomotion. We hypothesize that studying septins can complement the current model of amoeboid motility which depends on septins, yet *via* an unknown mechanism (Tooley et al., 2009). Previous reports show that septins participate in the formation of contractile actomyosin rings during cytokinesis, indicating a possible link between septins and amoeboid migration (Gilden et al., 2012; Tooley et al., 2009). Particularly, the septins’ GTPase activity drives the assembly of major septin isoforms (*i.e.,* SEPT2, SEPT6, SEPT7, and SEPT9) into non-polar filaments (Spiliotis, 2018; Woods and Gladfelter, 2021) that stabilize actomyosin structures, such as stress-fibers (Dolat et al., 2014; Schmidt and Nichols, 2004; Spiliotis and Nakos, 2021) and cytokinesis rings (Founounou et al., 2013; Kinoshita et al., 2002; Mostowy and Cossart, 2012). It is proposed that septins-based stabilization of actomyosin structures ensures the accumulation of substantial contractile energy (Dolat et al., 2014; Schmidt and Nichols, 2004), which facilitates both the dynamics of the amoeboid cortex in T cells (Gilden et al., 2012; Tooley et al., 2009) and the cortex-driven cytokinesis (Estey et al., 2010). Therefore, we decide to study whether septins participate in the amoeboid motility of T cells, *i.e.*, by regulating either actomyosin contractility or non-adhesive cell-microenvironment interactions within a structurally discrete microenvironment such as a 3D collagen gel.

We show that septins provide extracellular matrix-guided assembly of F-actin and alpha-actinin-rich cortical rings in migrating T cells. These rings compartmentalize the contractile actomyosin cortex into peristaltic treadmilling ‘ball-chain’ of spheroid segments. The lined-up cell segments feature ‘hourglass’ topography and discrete actomyosin contractility of individual segments, which facilitates transient steric stabilization of cells on obstacles and obstacle-evasive peristaltic translocation of cytosolic content between the adjacent cell segments. Consequently, we show that acute inhibition of septins’ GTPase activity causes an abrupt loss of cell segmentation along with the obstacle-evasive 3D propulsion. Our results indicate that the amoeboid motility of immune cells is a non-stochastic 3D circumnavigation guided by the structure of a crowding microenvironment and septin-based partitioning of actomyosin contractility.

## RESULTS

### Human CD4+ T cells subdivide cell cortex during amoeboid migration

Our principal observation is that primary human T cells (*e.g.,* human CD4+ T cell, hCD4+) compartmentalize the actomyosin cortex during amoeboid migration within 3D collagen-I gels. The typical migrating T cell changes its shape from a singular spheroid into a ‘ball chain’-like system of discrete and interconnected spheroid segments **(Figure 1a-1)**. These segments are separated by cortex-indenting circumferential furrows, identified as cortical F-actin rings **(Figures 1a-3**, **1b,** *F-actin, arrows*). The lined-up segments visually conform to available lumens within the surrounding collagen matrix and constrict the translocating nucleus **(Figures 1a-2**, **1c**, *nucleus, arrows*).

**Figure 1.**
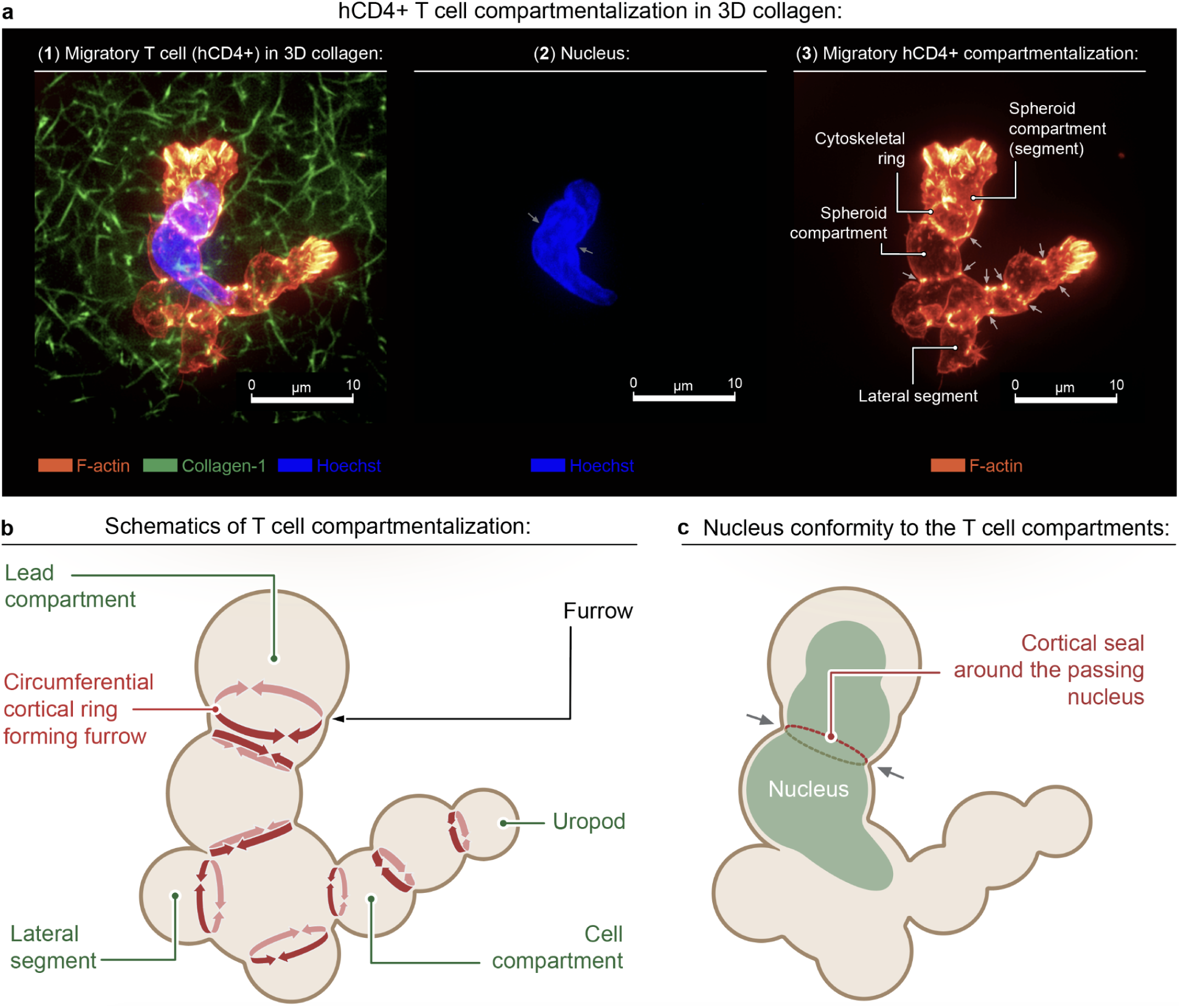
Compartmentalization of T cells during amoeboid migration in collagen gels. **(a)** Compartmentalization of the human CD4+ (*i.e.*, hCD4+) T cell cortex into multiple spheroid segments during amoeboid migration through a fibrous 3D collagen matrix (collagen type-I). Spheroid segments are separated by F-actin-enriched circumferential rings (SiR-actin staining for F-actin) that shape cortex (*arrows*) with the multiple circumferential furrows and indent a passing-through nucleus (*arrows*). **(b)** Schematics of T cell compartmentalization. Newly formed cortical F-actin ring (circumferential furrow) separates an expanding cell segment (leading compartment) from the previously formed segments within the trailing ‘ball chain’-shaped cell body.

The segmented morphology of migrating T cells, *i.e.,* the lined-up segments separated by discrete F-actin rings, reminisces the septin-stabilized membranous and/or cortical rings, reported in the dividing yeasts (Oh and Bi, 2011) and some mammalian cells (Bridges and Gladfelter, 2015; Founounou et al., 2013). Notably, we identify septins as a structural component in the leading F-actin ring in segmented T cells (*e.g.,* septin-7, **Figures 2a-c**, **+DMSO)**. Previously, structural and migration signatures of septins in immune cells have been studied only in 2D locomotion systems, with no specific role for septins in the motility of lymphocytes suggested to date (Gilden et al., 2012; Tooley et al., 2009). For example, only a punctuated pattern of septin striation has been observed in the T cells crawling on the flat surfaces during their 2D ‘random-walk’ motility, while a knockdown of septin-7 resulted in a T cell uropod elongation and decreased ‘random-walk’ efficiency (Tooley et al., 2009). Therefore, the identification of septin-positive cortical rings in migrating T cells could provide a new structural link between the reported correlation of septins’ expression (*e.g.,* septin-7) and the amoeboid behavior of T cells (Tooley et al., 2009).

**Figure 2.**
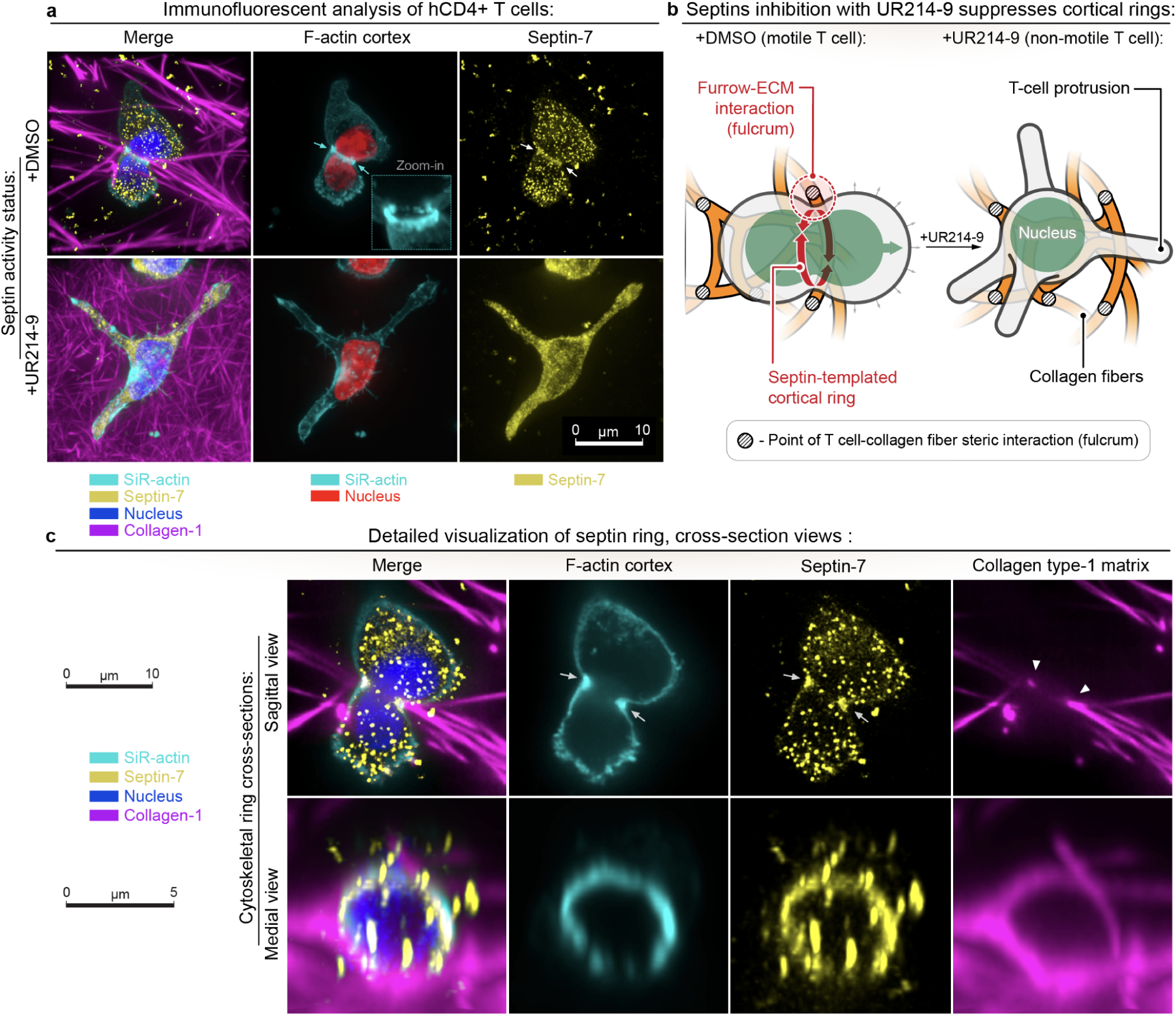
Septins GTPase activity simulates compartmentalization of T cells during amoeboid migration in collagen gels. **(a)** Immunofluorescence analysis of T cell migration in 3D collagen matrix: *Top* - hCD4+ T cell actively migrates in control conditions **(+DMSO)**, while displaying amoeboid compartmentalization with a septin-positive F-actin ring (Septin-7 immunofluorescence and SiR-actin staining, *arrows*). The migrating amoeboid T cell translocates its deformable nucleus through the cortical ring (*arrows*). *Note that in control T cells, immunofluorescence staining shows septin-7-enriched cytosolic vesicles and cortical structures*. *Bottom* - Migrating hCD4+ T cell loses amoeboid segmentation and the septin-positive cortical F-actin rings upon septin inhibition **(+UR214-9)**. Instead, the T cell acquires a non-migratory cell morphology, which is characterized by the formation of multiple elongated linear protrusions. *Note that UR214-9 also causes the loss of discrete septin-enriched cytosolic vesicles and cortical structures (as shown by a diffuse septin-7 immunofluorescence staining)*. **(b)** Schematic representation of how an amoeboid T cell transitions to the non-migratory phenotype. *Left* - An amoeboid T cell in the sterically complex microenvironments, *e.g.*, collagen matrix, forms a cortical ring (sterically interactive furrow) between the adjacent spheroid compartments of larger diameters **(+DMSO)**. During the amoeboid propulsion in 3D collagen, spatial configuration of adjacent segments provides a transient steric lock (fulcrum) between the cell and confining collagen fibers. The fulcrum mechanically supports peristaltic translocation of cytoplasm and nucleus without friction-loaded displacement of the entire T cell cortex. *Right* - A T cell loses the amoeboid migration and cortical response (cortical F-actin ring) to the sterically complex microenvironment upon chemical suppression of septins’ GTPase activity with UR214-9. **(c)** Detailed transverse and sagittal cross-section views of the septin-enriched (septin-7 immunofluorescence) cortical rings in the migrating T cell (see **c**, *top*). *Note that the septin-positive cortical F-actin ring forms inside the confining collagen lumen (arrows)*.

To study the role of septins in the formation of cortical rings, we proceed with inhibiting septins’ GTPase activity with a novel mammalian septins’ GTPase activity inhibitor, UR214-9 (Kim et al., 2020; Zhovmer et al., 2021). The acute chemical perturbation of septins causes an abrupt loss of the amoeboid organization in hCD4+ T cells (**Figure 2a**, **+UR214-9,** 40 µM, 1 hour), which manifests as **(1)** loss of cell segmentation, **(2)** loss of cortical rings, and (**3)** transition to non-migratory morphology, characterized by the passive cell shape conformity to the surrounding collagen matrix, i.e., *via* multiple elongated cell protrusions. The loss of T cell segmentation and simultaneous transitioning between the migratory **(Figure 2b, +DMSO)** and non-migratory phenotypes **(Figure 2b, +UR214-9)** suggest that the ‘hourglass’-shaped cell surface formed by a cortex-indenting circumferential furrow (*i.e.,* cortical ring) and two adjacent spheroid cell segments may actively participate in the amoeboid motility, forming steric support (fulcrum) by transiently latching the cell cortex onto confining collagen fibers (**Figure 2b**, **+DMSO)**.

### Amoeboid migration requires actomyosin and septins GTPase activity

To confirm the participation of septins in 3D amoeboid motility, we quantify hCD4+ T cell migration in the tissue-like 3D collagen matrix (**Figure 3a**) while perturbing septin’s GTPase activity. As described above, static images of T cells indicate that they robustly segment their actomyosin cortex by cortical rings during migration in 3D collagen matrices (**Figures 1-2**, **+DMSO)**. Indeed, the control T cells demonstrate robust treadmilling of cell segments (**Movie 1**) and a stepwise segment-to-segment translocation of cytoplasm and nucleus through the cortical rings **(Movies 2** and **3**, **Figure SI1)**. Notably, septins inhibition substantially reduces the speed of migrating T cells to the near arrest (**Figure 3b**, **+DMSO** *vs.* **+UR214-9)** and substantially decreases their effective migratory displacement (**Figure 3c**, **+DMSO** *vs.* **+UR214-9)**.

**Figure 3.**
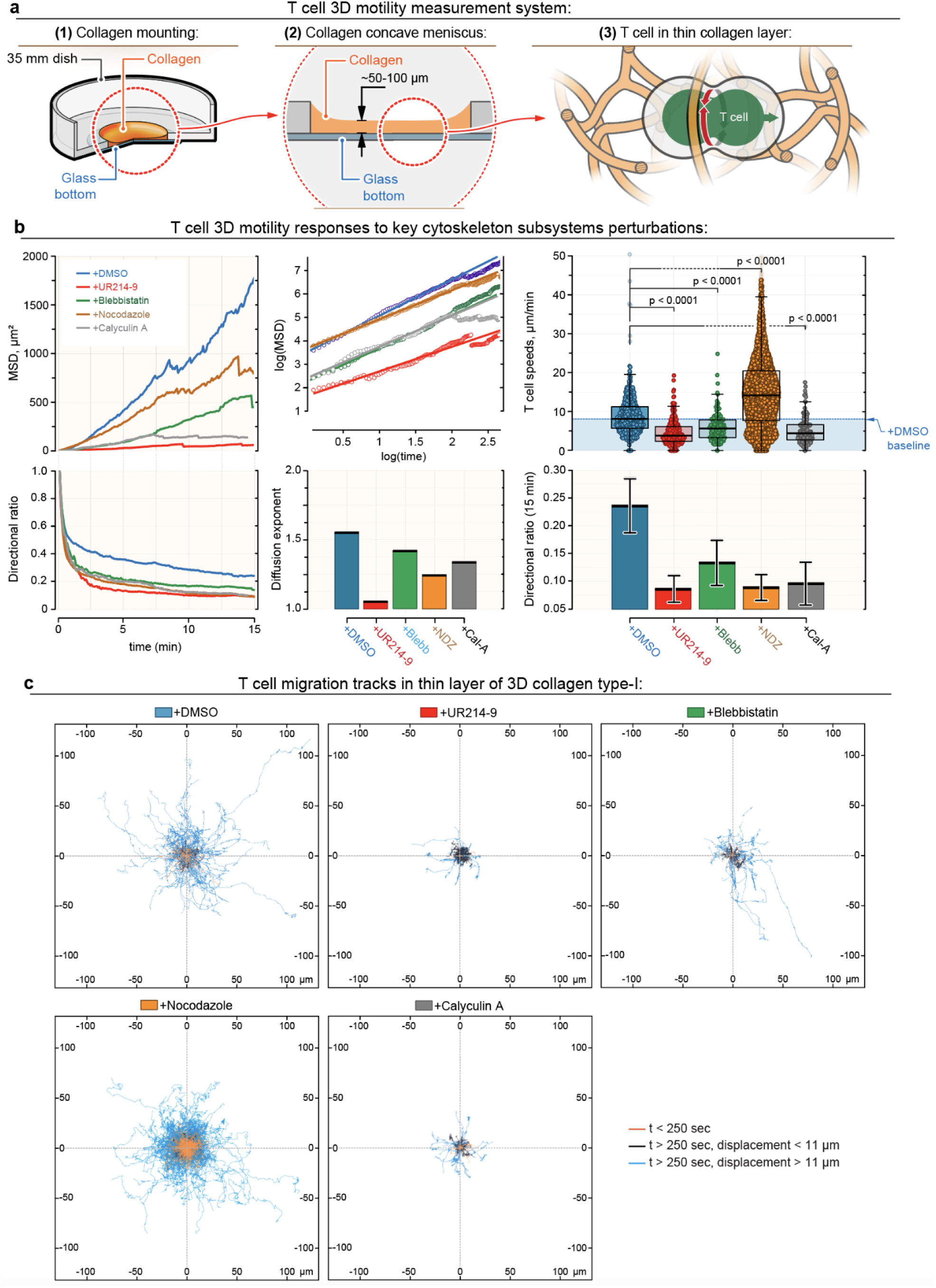
Perturbation of septins’ GTPase activity arrests amoeboid migration of T cells within the 3D microenvironment. **(a)** Schematic representation of live cell imaging platform (**1**) used to measure motility of human CD4+ T cells in the 3D collagen matrix. Migrating T cells are imaged within a thin layer of collagen gel (**2**) and visualized in the planes positioned away from the glass (**3**). **(b)** Responses of the characteristic motility parameters for human CD4+ T cells migrating within a 3D collagen matrix during perturbation of the key cytoskeletal subsystems: septins **(+UR214-9)**, microtubules **(+Nocodazole**, *i.e.* +NDZ), and non-muscle myosin contractility **(+Blebbistatin**, *i.e.* +Blebb, and **+Calyculin A**, *i.e.* +Cal-A). *Note that mean velocity values of automatically detected and tracked T cells do not reach zero due to the lack of sufficient T cells-collagen adhesion. Insufficient adhesion enables lymphocytes’ passive oscillations (e.g. Brownian motion) within a 3D collagen matrix, which includes completely arrested T cells (e.g.* ***+UR214-9*** *or* ***+Calyculin A*** *treatments). Passive T cell oscillations do not result in the lymphocyte’s persistent displacement within the collagen matrix*. MSD - Mean Square Displacement is the population-wide average of the squared values of the linear distance between the T cell position at time *t* and its initial position on the track (*t* = 0). Displacement ratio is the population-wide average value of the ratios of the linear distance between the T cell position at time *t* and its initial position on the track to the distance that T cell has traveled along its curved track up to time *t*. Note that a cell moving strictly along a line would have the Displacement ratio of one, while a cell wandering around its initial location would have the Displacement ratio close to zero. Thus, Displacement ratio is a metric of directional persistence. Diffusion exponent is the slope of the linear fit to MSD in log-log scale. Note that for a particle undergoing Brownian motion in 3D, MSD = 6*Dt*, where *D* is the diffusion constant. More generally, MSD can be approximated as MSD∼*t^a^*, so that in log-log scale, we have approximately linear dependence: log(MSD) = *a* log(*t*) + constant, where *a* is the Diffusion Exponent. Thus, the Diffusion Exponent is a metric of the deviation of cell trajectories from the normal Brownian motion (superdiffusion/active transport for *a*>1 and subdiffusion for *a*<1). **(c)** 2D projections of the human CD4+ T cell migration tracks within a thin layer of 3D collagen matrix during perturbation of the key cytoskeletal subsystems. *Note that both* ***+UR214-9*** *and* ***+Calyculin A*** *treatments induce potent arrest of T cell displacement, due to the complete loss of amoeboid cortical dynamics*.

Previous reports indicated that actomyosin contractility is important for amoeboid dynamics in T cells (Tabdanov et al., 2021; Takesono et al., 2010). Consequently, our data confirm that contractility inhibition **(+Blebbistatin,** 25 µM) reduces T cell speed and displacement (**Figure 3b-c**, **+DMSO** *vs.* **+Blebbistatin,** 1 hour), although with lower efficiency than UR214-9 (**Figure 3b-c**, **+UR214-9** *vs.* **+Blebbistatin)**.

The stimulation of the actomyosin contractility *via* MT→GEF-H1→RhoA signaling axis, achieved by microtubule depolymerization **(+Nocodazole**, 5 µM) (Krendel et al., 2002; Tabdanov et al., 2021; Takesono et al., 2010), substantially increases migration speed (**Figure 3b**, **+Nocodazole**, 1 hour), but not the effective 3D displacement (**Figure 3c**, **+Nocodazole)** due to the reduced directionality persistence of cell migration (**Figure 3b**, **‘*diffusion exponent’*** and **‘*directional ratio’*)**.

On the contrary, the activation of non-dynamic actomyosin contractility, *i.e.,* by locking actomyosin in the hypercontractile state with high tension and low dynamics, collapses T cells into immotile spheroids (**Figure 3b-c**, **+Calyculin A**, 50 nM, <1 hour). These results indicate that the amoeboid motility of T cells primarily depends on the dynamic actomyosin contractility and septin GTPase activity.

### Septins work in conjunction with actomyosin

To further investigate how the amoeboid motility may depend on the actomyosin and septins, we use a strategy of acute pharmacological inhibition of septins GTPase activity. This strategy potentially decreases the probability of causing complex, non-linear, and poorly decipherable cumulative downstream effects compared to available alternatives, such as long-term perturbation of septin activity by genetic knockout. The applied septin inhibitor UR214-9 is a novel fluoride-derived analog of a better-known fungal (*i.e.,* yeast) septin inhibitor - forchlorfenuron (FCF) (Hu et al., 2008), which, unlike FCF, displays negligible cytotoxicity towards mammalian cells at high-doses (Kim et al., 2020; Singh et al., 2020; Zhovmer et al., 2021).

To test that the UR214-9 acts like FCF, *i.e.,* provides an abrupt and detectable loss of septin-actomyosin interactions (Cheffings et al., 2016; Dolat et al., 2014; Founounou et al., 2013; Salameh et al., 2021; Schmidt and Nichols, 2004), we choose the mesenchymal triple negative adenocarcinoma MDA-MB-231 cell line. Compared to T cells, MDA-MB-231 cells feature distinct contractile actomyosin structures, *i.e.,* stress-fibers. Using imaging of stress fibers, we visualize that acute inhibition of septins GTPase activity **(+UR214-9)** results in rapid translocation of septins (*e.g.,* endogenous septin-9, immunofluorescent analysis) from the stress fibers to the cytosol **(Figure SI2a**, **+DMSO** *vs.* **+UR214-9)**. Rapid translocation of septins is accompanied by the structural dissolution of stress fibers resulting in partial cell retraction and detachment **(Figure SI2a**, **+UR214-9**, 40 µM, 1 hour). Similarly, in live cells transfected with GFP-septin-9, the collapse and disintegration of septin-positive filaments into the curled fragments **(Figure SI2b**, **+DMSO**, **+UR214-9**, 1 hour) occur during the first hour after the addition of UR214-9.

In the orthogonal experiment with MDA-MB-231 cells, we suppress actomyosin contractility by targeted myosin II inhibition with Blebbistatin which results in a similar collapse of the stress-fibers decorated with septin-9, as shown by fragmentation and redistribution of GFP-septin-9 filaments that are no longer associated with stress fibers **(Figure SI3c**, **+Blebbistatin**, 50 µM, 1 hour). These results suggest that septins and contractile actomyosin structures are mutually interdependent, *i.e.,* septins stabilize the tensile actomyosin, while loss of myosin-driven F-actin contractility destabilizes the associated septin structures, supporting previous findings that homozygous loss of septin-9 in mouse embryonic fibroblasts causes a loss of stress fibers and associated septin filaments (Füchtbauer et al., 2011). Moreover, our experiments indicate that UR214-9 acts like FCF, causing the loss of septin-actomyosin interactions.

### Septins sense cortex-indenting obstacles

To study the role of septin-actomyosin interactions in 3D motility, we analyze where T cells typically feature distinct septin-enriched cortical rings. We note that these rings are found exclusively at the cell surface sites indented by collagen fibers (**Figure 4a**, *arrows*). Analysis shows that control T cells **(+DMSO)** have an average of 1 to 2 rings per cell (**Figure 4b**, *left*), and no cortical rings are observed without a cell surface-indenting contact with collagen fibers (**Figure 4b**, *right*).

**Figure 4.**
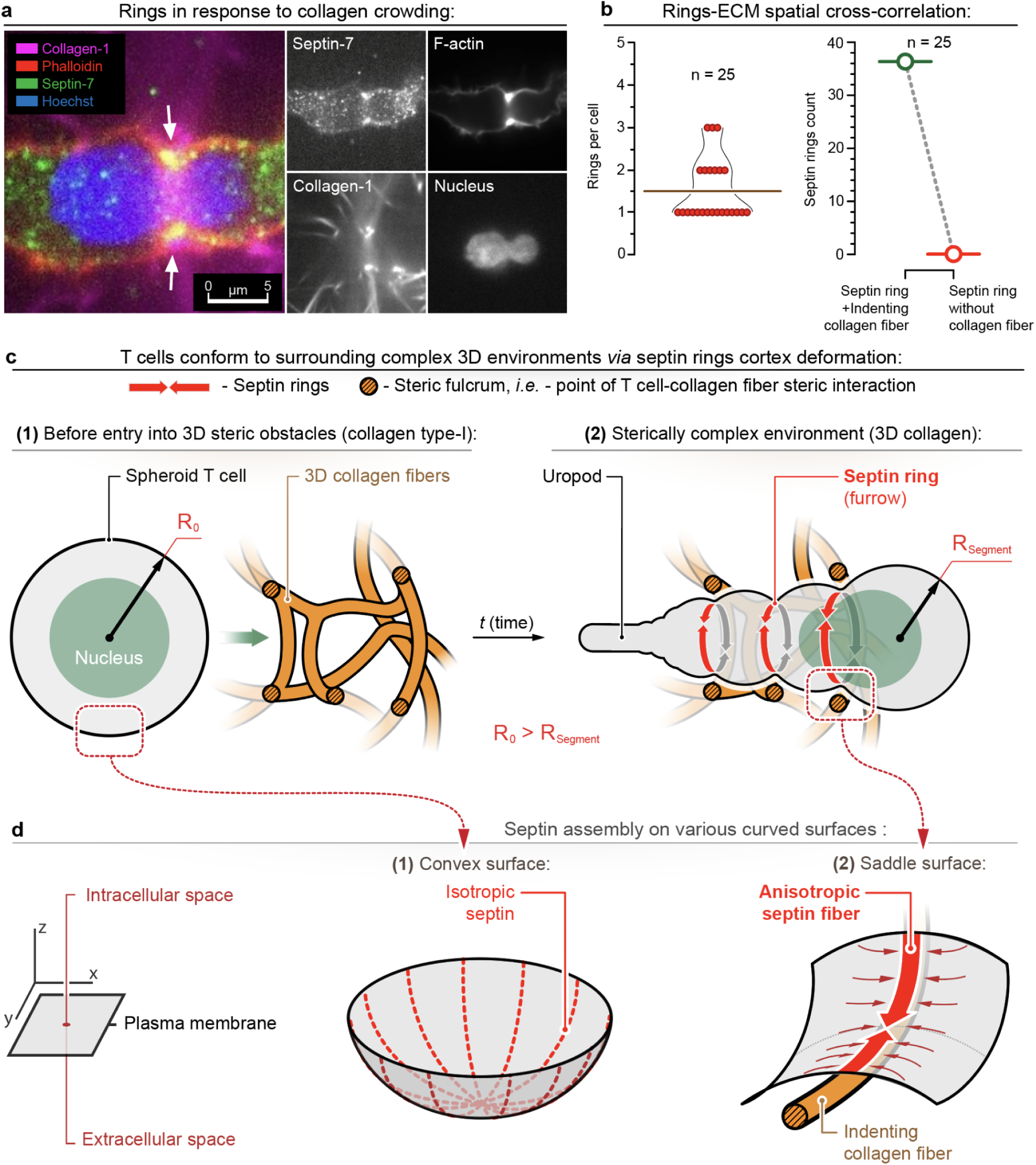
3D structure of the extracellular matrix guides the formation of septin-positive cortical rings. **(a)** Sagittal cross-section views of the amoeboid T cell migrating through the 3D collagen matrix. *Note that the septin-positive (septin-7) cortical ring forms at the site of the T-cell mechanical interaction (cell surface-indenting) with the collagen fibers (arrows).* The formation of the cortical ring at the indentation sites allows for cell compartmentalization and conformational adaptation to the complex configuration of the surrounding obstacles, as well as for the peristaltic translocation of the cytoplasm and nucleus along the sequence series of the resulting cell segments. *Note that the nucleus conforms to the surrounding cortical ring and collagen fibers*. **(b)** Immunofluorescent analysis of spatial colocalization and cross-correlation for the cortical rings and cell surface-indenting collagen fibers. *Left* - Distribution of the rings-per-cell counts (mean value ∼1.5 ring/cell). *Right* - Cortical rings are linked to the presence of cell surface-indenting ECM (collagen) fibers. *Note that rings are always associated with the adjacent collagen fiber (ring+collagen), while there is no identified T cell ring without a presence of a T-cell-indenting collagen fiber*. **(c)** Schematic representation of the ECM-induced T cell transitioning from the non-migratory spheroid to the migratory compartmentalized architecture during its amoeboid migration through the sterically interactive 3D collagen gel. External 3D topography of collagen fibers meshwork surrounding an initially spheroid T cell **(1)** provides a guiding spatial configuration of the steric and mechanical pressure stimuli, to initiate a series of septin-driven T-cell compartmentalization events (**2**) required for amoeboid migration. Note that the resulting T cell’s segment radii (R_Segment_) are smaller than the radius (R_0_) of the spheroid T cell. **(d)** Schematic representation of septins assembly on the inner side of the T-cell plasma membrane surfaces with various curvatures. (**1**) - isotropic assembly of septin filaments on convex surface, *e.g.* in a spheroid cell. (**2**) - anisotropic assembly of septins on the saddle surface formed by a cell surface-indenting collagen fiber, *e.g.* initiation of the assembly of the compartmentalizing septin rings in a migrating T cell in response to the external obstacles.

From the mechanistic standpoint, septins are well-known sensors of the plasma membrane curvatures (*i.e.,* second-order surface curves) that preferentially assemble into isotropic networks on the inside of spheroid (convex) surfaces, avoid concave surfaces, and self-organize into the macroscopic fibers along the spine of the indenting furrow on the inside (mixed, *i.e.,* - saddle) surfaces of the cell membrane (Beber et al., 2019) (**Figure 4c-d**). For example, we observe curvature-sensitive accumulation of septin-7 (SEP7-GFP) filaments along the convex ridge in the MDA-MB-231 cells (**Figures 5a-b**, SI3).

**Figure 5.**
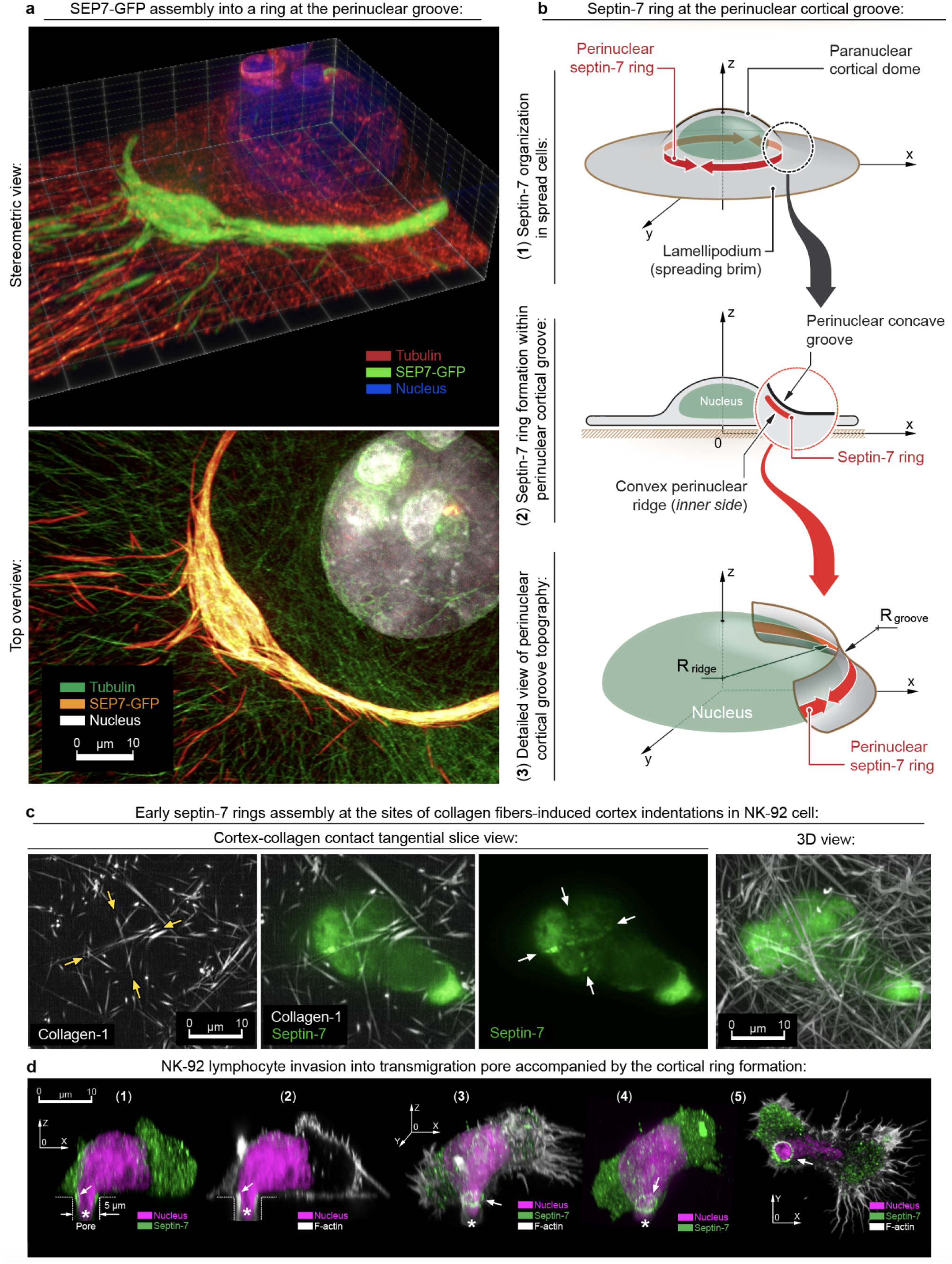
Membrane (cortex) curvature-sensitive accumulation of septin-7 (SEP7-GFP) filaments at the inner side of deflection, *i.e.*, along the convex ridge in the MDA-MB-231 cells or collagen fiber-induced indentation in NK-92 lymphocytes. **(a)** Detailed iSIM microscopy view of the perinuclear region that features a deflection of the cell cortex/plasma membrane surface around the nucleus-induced cell dome, *i.e.*, perinuclear concave groove (convex ridge on the inner side of the cortex). *Note a selective accumulation of the SEP7-GFP filaments at deflection regions of the cell plasma membrane and cortex*. **(b)** Schematic representation of curvo-sensitive behavior of SEP7, forming the filaments aligned along the convex ridge at the perinuclear region. **(c)** Tangential slice views of the amoeboid NK cell (NK-92 cell line) migrating through the collagen matrix. Early stage of septin-7 assembly (Septin-7 immunofluorescence, *green*) into the cortical ring starts at the sites of T cell surface indentation, caused by the T cell pressure against the collagen fibers *(arrows)*. **(d)** 3D views of a NK-92 lymphocyte entering the 5 µm transmigration pore (*dashed lines*) with an invading segment (*asterisk*). (**1-2**) - Sagittal cross-section view indicates assembly of septin-7 (**1**) and F-actin (**2**) dense cortical structure at the entrance into the pore (*arrows*). (**3**) - Stereometric 3D views of the NK-92 cell shows a complete and mature cortical ring (*arrow*). (**4**) - Detailed septin-7 ring view (*arrow*) at the neck of the invading segment. (**5**) - View from the bottom at the invading protrusion (*arrow*) NK-92 cells.

Thus, a possible scenario of septin-7 assembly is that in T cells, collagen fibers, by crowding interactions with the membranes of migrating T cells (Wolf et al., 2003), create cortex-indenting mechanical collisions between the cell cortex and surrounding ECM obstacles with a curvature that is sufficient to trigger the local assembly of septins (**Figure 4c**). Moreover, our observations indicate that curvature-guided septin assembly may happen across various types of immune cells. For example, natural killer NK-92 lymphocytes are able to condense septin fibers along the collagen fibers-induced cortex indentations (**Figure 5c**, *arrows*). Similarly, NK-92 lymphocytes, passing through 5 µm plastic pores, form the septin-7-rich and F-actin-dense circular structures (**Figure 5d**), which resemble cortical rings and septin-mediated plant invasion structures described for fungi (Dagdas et al., 2012).

### Septins compartmentalize actomyosin contractility for circumnavigation

For T cells and other lymphocytes, the microenvironment-guided cell compartmentalization was previously unrecognized. Structurally, this cell compartmentalization by cortex-indenting collagen fiber manifests as a sterically interactive furrow marked by the assembly of a septin-enriched cortical ring (**Figure 6a**). We propose that the ‘segment-ring-segment’ topographic unit of gel-invading T cell mechanically latches its cortex on collagen and serves as a non-adhesive, *i.e.,* steric, fulcrum that facilitates peristaltic translocation of the nucleus and cytosolic content into the newly formed invading cell segment (**Figure 6b**). This stepwise peristaltic translocation of the nucleus and cytosolic content along the treadmilling chain of forming and shrinking T cell segments would require spatiotemporally coordinated contractility of the actomyosin cortex to provide directed cytosolic hydrostatic pressure (Dolat et al., 2014; Salameh et al., 2021). Indeed, immunofluorescence analysis of segmented T cells reveals an uneven spatial distribution of activated phosphorylated (Ser19) myosin across adjacent cell cortical segments **(Figure 6c-1)**. The differential density of activated cortical actomyosin between individual segments **(Figure 6c-2)** indicates that each cell segment can feature an individual level of cortical contractility required for spatiotemporally coordinated peristaltic dynamics of the entire cell cortex.

**Figure 6.**
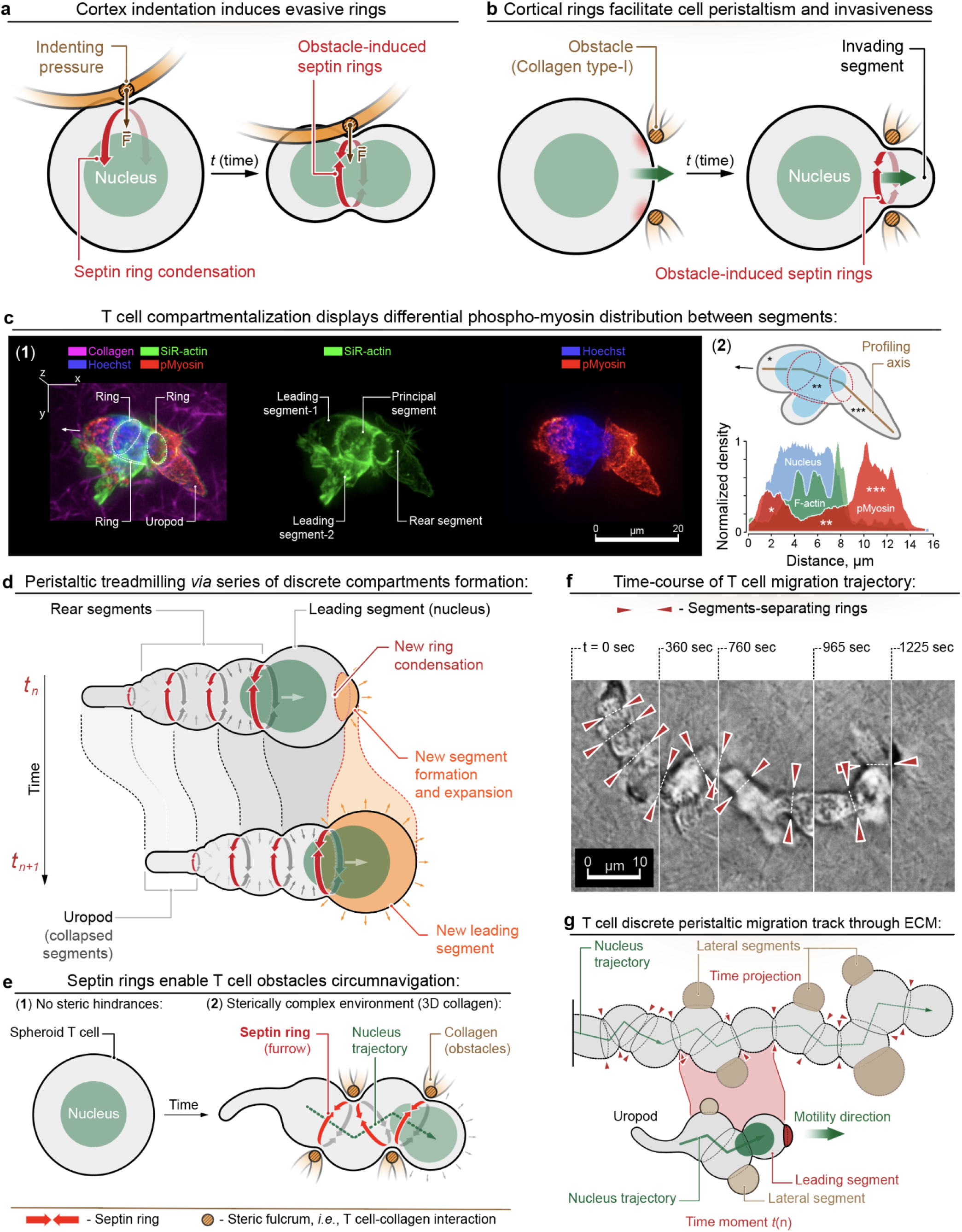
Actomyosin cortex compartmentalization by localized formation of the septin rings in response to the external obstacles enables T cell peristaltic circumnavigation within complex 3D microenvironments. **(a)** Schematic representation of evasive cortex deformation, *i.e.* cortical ring, initiated by a cell surface-indenting collagen fibers. The spatial configuration of the resulting furrow forms a transient steric lock, *i.e.* fulcrum, onto indenting collagen fibers by virtue of cortical ring-induced ‘hourglass’ cell topography. **(b)** Schematic representation of adhesion-independent invasive amoeboid propulsion. Collagen-induced cell partitioning initiates a stepwise peristaltic translocation of cytosolic content into the invading cell segment without friction-loaded displacement of the entire T cell cortex. **(c)** (**1**) - 3D micrographic view of the hCD4+ T cell **(+DMSO)** in the collagen matrix. The T cell forms multiple rings and discrete contractile cell segments. *Note that the cortex in adjacent segments displays a differential density of phospho-myosin (phospho-myosin light chain 2 (Ser19) immunofluorescence), indicating that each individual cell segment may feature a different level of cortical contractility.* (**2**) - Cumulative density profiling for phospho-myosin light chain 2, F-actin and chromatin along the T cell’s main axis, *i.e.* - across the principal segments. *Note that phospho-myosin peaks in the leading and the rear segments, but is depleted in the main (middle) nucleus-containing cell segment*. **(d)** Schematic representation of amoeboid T cell migration by peristaltic treadmilling of discrete cell segments. Series of the new compartments formation at the leading front of the cell is accompanied by the nucleus and cytosol translocation **(Movies 1, 2 and 3)**. **(e)** Schematic representation of T cell circumnavigation during amoeboid migration. Collagen fibers-induced septin rings deform the T cell cortex to conform to the surrounding obstacles. Treadmilling of the compartments and cytosol/nucleus peristaltic propulsion throughout the series of the obstacles-evading cortical rings allows T cells to efficiently circumnavigate around and between the structurally complex collagen matrix. **(f)** Time course collage of the single hCD4+ T cell along its migration track within the 3D collagen matrix. Note the T cell forms multiple segments (chambers), separated by the rings (arrowheads) along the track of its migration. The T cell trajectory is a zigzag track between the chambers that are formed during the migration. **(g)** Schematic representation of the spatiotemporal superposition of the migrating T cell - motility is driven by peristaltic translocation of the nucleus between segments, separated by the cortical rings (highlighted by arrowheads). Nucleus translocation forms the main zigzag track that reflects the directionality of the new leading segment formation, guided by the surrounding microenvironment steric configuration and topography. *Note that T cells also develop side branching pseudopods (lateral segments) that do not accommodate the nucleus and therefore do not become the leading segments*.

Our immunofluorescence data also aligns with the live dynamics of translocation of cytosol and nucleus across multiple segments during 3D migration of T cells in the collagen matrix **(Movies 1**, **2,** and **3**, **Figure SI1)**. In particular, we observe that each locomotion cycle in a migrating T cell includes propulsion of cytosol and nucleus and initiation of a new cell segment. The ‘old’ or utilized cortical segments, which are located behind the nucleus, undergo gradual degradation, recycling, and compaction, manifesting as the rear-located cylindrically shaped uropod (**Figure 6d**).

Based on the existing model for the stochastic cortical instability as a driving force of amoeboid motility (Ruprecht et al., 2015), we suggest a model of steric guidance, which is a non-stochastic cortical partitioning in amoeboid cells driven by ECM structure. Specifically, we suggest that the mechanical collision between the expanding cell segment and the confining steric obstacles creates cortical indentation, which may trigger septin condensation and assembly of a new cortical ring with the separation of a new cell segment. The treadmilling of ECM-conformed cell segments provides lymphocytes with a mechanism for a stepwise obstacle-avoiding peristaltic translocation of cytosol and nucleus between cell segments (**Figure 6e**).

For a complex tissue-like environment, *e.g.,* collagen matrix, such stepwise ECM-guided peristaltic motility manifests as circumnavigation, *i.e.,* the obstacle-evasive zigzag-like trajectory of amoeboid cells. Notably, live imaging of the multi-segmented hCD4+ T cells within the 3D collagen matrix shows that lymphocytes repeatedly form a zigzagging trajectory (**Figure 6f-g**, **Movie 1)**, emphasizing the migration path between and around collagen fibers through the available lumens (*i.e.,* circumnavigation). During circumnavigation, T cells continually treadmill old shrinking and newly-formed expanding discrete cortical segments formed in response to cell surface-indenting steric obstacles (**Figure 6g**).

Additionally, we identify that cortical rings are enriched in alpha-actinin (**Figure 7a-b**) and depleted in myosin 2a (**Figure 7c**). These results indicate that heavily cross-linked cortical F-actin rings in T cells are likely more similar to non-contractile and non-stretchable solid-like actomyosin structures (Lieleg et al., 2009), which may mechanically stabilize the expanding cell cortex against the applied external stimuli, hence, locally reducing its excessive hydrostatic pressure against the indenting obstacles and creating the obstacles evasion event (**Figure 7e**).

**Figure 7.**
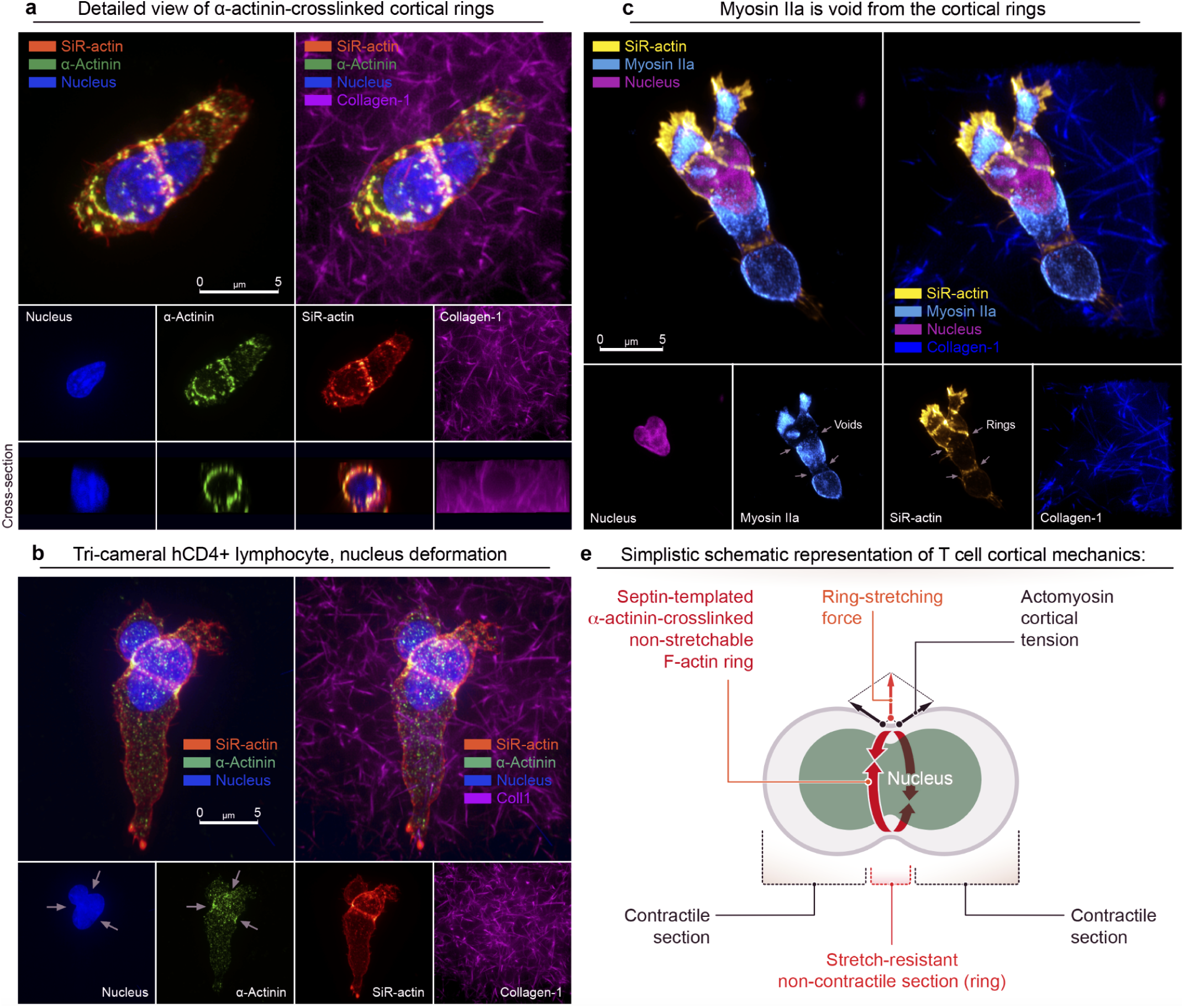
Septin-templated α-actinin-crosslinked cortical rings are not contractile, but provide the non-stretchable mechanics for efficient T cell cortex’ compartmentalization. **(a)** Detailed view of the cortical rings reveals α-actinin as cortical rings’ F-actin’s crosslinker (periodic structures). **(b)** Visualization of the non-stretchable mechanics of the α-actinin-crosslinked F-actin cortical rings (nucleus deformation, *arrows*). **(c)** Myosin IIa is depleted from the α-actinin-crosslinked cortical F-actin rings (*arrows*), indicating their non-contractile, yet stretch-resistant mechanics. **(e)** Schematic representation of the lymphocyte’s cortical mechanics: myosin IIa-enriched (*i.e.*, contractile) spheroid segments are separated by the stretch-resistant, yet non-contractile α-actinin-crosslinked F-actin rings.

## Discussion

We show that primary human CD4+ T cells compartmentalize their actomyosin cortex during amoeboid migration in response to steric cues, as demonstrated by the ECM-guided formation of cortical rings. Moreover, we identify septins in these rings and demonstrate that inhibition of septins’ GTPase activity causes the abrupt loss of cortical rings and cell segmentation as well as transitioning to non-migratory morphology. These data suggest a non-stochastic mechanism of amoeboid motility for immune cells that is alternative yet complementary to both the previously described mechanism of amoeboid motility by blebbing and the non-adhesive sliding motility in non-immune melanoma cells (Adams et al., 2021; Logue et al., 2015).

Briefly, the basis for sliding motility in amoeboid melanoma cells is a cortex treadmilling within the lasting leader bleb that powers friction-based cell displacement within an ideal “sandwich”-like microenvironment, *i.e.,* between two confining parallel flat surfaces (Adams et al., 2018; Liu et al., 2015). Although the described sliding is considered a particular example of the wide and poorly characterized variety of amoeboid motility mechanisms, the locomotion based on friction seems energetically and mechanically suboptimal. Therefore, we question whether the observed compartmentalization of T cells in collagen gels enables a more optimized locomotion mechanism inherent to immune cells in tissues. We propose that the ECM-guided subdivision of the whole cell into a series of relatively independent contractile segments and their discrete treadmilling allow hindrance-evasive peristaltic motility without friction-loaded displacement of the entire T cell cortex against a structurally crowded 3D microenvironment.

Our conjecture is based on the observation of amoeboid sliding motility in non-immune melanoma cells, where an excessive cross-linking of contractile cortical F-actin (*e.g.,* by filamin-A and/or fascin), combined with the cells’ sterical confinement, results with cortex reorganization into a continuous system of coaxial circumferential fibers, forming an elongated cylindrical cortex of a leading bleb (Adams et al., 2021). We suggest that the reported filamin-A and/or fascin-rich circumferential cortical structures in melanoma cells and the observed discrete septin-positive cortical rings in human T cells are both analogous to the linear stress-fibers. Moreover, both cortical structures and stress fibers depend, at least in part, on the scaffolding and stabilizing effects of septins (Dolat et al., 2014; Kremer et al., 2007; Salameh et al., 2021).

To support this idea, we demonstrate that septins may act as mechanosensitive F-actin cross-linkers, preferentially binding and stabilizing tensile actomyosin structures, as demonstrated by rapid dissociation of stress fibers and translocation of septins from actomyosin upon acute inhibition of either actomyosin contractility or septins GTPase activity. Therefore, similarly to stress-fibers that condense from an isotropic actomyosin network into a load-resisting actomyosin band between anchoring focal adhesions (Lehtimäki et al., 2021; Vignaud et al., 2021), T cells may use septins to facilitate condensation of F-actin and alpha-actinin-rich cortical rings as a circularized (*i.e.,* anchorless) form of the stress-fiber in response to cortex indentation. Moreover, we propose that crosslinker-heavy but myosin-depleted cortical rings of T lymphocytes are non-contractile by design, preventing potential cytokinesis-like abscission of spheroid cell segments. Instead, non-stretchable cortical rings serve to resist further stretching forces provided by myosin-rich contractile spheroid cell segments, *e.g.,* by limiting unsafe cortex-expanding hydrostatic pressure against surrounding steric obstacles. Thus, while mesenchymal cells use anchored myosin-rich stress fibers to enable the mechanosensing of the adhesive microenvironments, the non-adhesive amoeboid cells utilize the anchorless myosin-depleted cortical rings for safe circumnavigation, i.e., obstacle avoidance, during the non-adhesive motility in sterically complex environments (Charras and Paluch, 2008; Paluch et al., 2005; Reversat et al., 2020; van der Gucht et al., 2005; Wolf et al., 2003).

In summary, our results indicate that T cells use septins for ECM-guided partitioning of actomyosin contractility, which could explain the previously reported links between amoeboid motility of T cells and septin-7 genetic knockdown (Gilden et al., 2012; Tooley et al., 2009). Specifically, we propose that cortex-indenting collagen fibers trigger septin assembly within the actomyosin cortex, which restructures the spherical T cell into a hydrostatically unified system of expanding, static, and contracting cortical segments. Obstacle-evasive formation and treadmilling of cell segments provides continuous shape-changing dynamics and configures cortical forces (*i.e.,* individual contractility of cell segments) to the pre-existing spaces, locking cell on the confining collagen fibers and exerting forces sufficient enough for safe hindrance-evasive peristaltic propulsion of nucleus and cytoplasm through the series of narrow lumens in 3D microenvironments.

## METHODS

### Cell experiments

We maintained human MDA-MB-231 cells (ATCC® HTB-26™) in DMEM with 4.5 g/L D-glucose, L-glutamine,110 mg/L sodium pyruvate (Corning Cellgro®, Cat# 10013CV) and 10% heat-inactivated FBS (HyClone®, Cat# SH30071.03H) at 37°C in 5% CO2.

Primary human CD4+ T cells were isolated from commercially available whole human blood (STEMCELL Technologies Inc., USA, Cat# 70507.1) with EasySep Human CD4+ T Cell Isolation Kits (STEMCELL Technologies Inc., USA), activated and expanded in ImmunoCult-XF T Cell Expansion Medium (STEMCELL Technologies Inc., USA) with the addition of ImmunoCult Human CD3/CD28/CD2 T Cell Activator and Human Recombinant Interleukin 2 (IL-2, STEMCELL Technologies Inc., USA) as per STEMCELL Technologies Inc. commercial protocol, at 37°C in 5% CO2.

NK-92 (ATCC®, CRL-2407™) were cultured in the ImmunoCult-XF T Cell Expansion Medium (STEMCELL Technologies Inc., USA) with the addition of Human Recombinant Interleukin 2 (IL-2, STEMCELL Technologies Inc., USA).

Cells were treated in glass-bottom 35 mm Petri dishes (MatTek Corp., Cat# P35G-1.5-14-C) using dimethyl sulfoxide (Sigma, Cat# 472301), (-)-Blebbistatin enantiomer (Sigma, Cat# 203391), Nocodazole (Abcam, Cat# ab120630), Dynapyrazole A (Sigma, Cat# SML2127), Calyculin A (Sigma-Aldrich, Cat# 208851), and septin inhibitor UR214-9 (synthesized by Dr. Rakesh K. Singh) as indicated in the main text.

Transfection of MDA-MB-231 cells with pEGFP-C2_SEPT9 i1 (Addgene Cat# 71609) and Septin-7 (VectorBuilder, Cat# VB220525-1339dzw) was performed in Opti-MEM (Gibco, Cat# 31985-070) using SuperFect reagent (Qiagen, Cat# 301307) according to the manufacturer’s protocol.

### Polymerization of prestained collagen gels

To prepare 3 mg/ml collagen-I gel, we assembled a gel premix on ice in a prechilled Eppendorf tube. To 1 volume of CellAdhere™ type I bovine, 6mg/mL (STEMCELL Technologies Inc., USA) we added 8/10 volume of DMEM, 1/10 volume of 10x PBS, 1/20 volume of 1M HEPES, and 1/20 volume of 1M Atto 647 NHS Ester (Sigma-Aldrich), Alexa Fluor 568 ester (Molecular Probes, Cat# A20003) or Alexa Fluor 488 ester (Molecular Probes, Cat# A20000). A drop of premixed gel (∼50 µL) was spread immediately on a glass surface of a plasma-treated glass-bottom 35 mm Petri dish (MatTek Corp., Cat# P35G-1.5-14-C) with a pipette tip. During polymerization (room temperature, for overnight), gels were covered with 1mL of mineral oil (Sigma-Aldrich, Cat# M8410) to prevent evaporation of water. Before adding T cells, polymerized gels were rinsed with PBS to remove the unpolymerized gel components.

### Synthesis of UR214-9

Equimolar mixture of 3-amino-2-fluorobenzotrifluoride (Combi-Blocks, Cat#QA-4188) and 2,6-dichloro-4-isocyanatopyridine (Toronto Research Chemicals, cat#159178-03-7) were stirred and heated in anhydrous Toluene at 85°C overnight. The separated UR214-9 was filtered and dried under vacuum in a dessicator. A portion of UR214-9 was detected in the Toluene layer, which was concentrated and purified by thin-layer chromatography using ethyl acetate and hexane as eluents. The pure product band was scrapped off the glass plate and UR214-9 was stripped from the silica gel using DCM+MeOH through a sintered funnel. The solvent was evaporated using a Buchi rotary evaporator to obtain UR214-9 as an off-white powder. The structure was confirmed by X-ray crystallography.

### Super-resolution and epifluorescent microscopy

For imaging, fixed cell samples were immunostained as indicated in corresponding Figures. Super-resolution imaging was performed using an Instant structured illumination microscopy (iSIM) system (VisiTech Intl, Sunderland, UK) equipped with an Olympus UPLAPO-HR ✕100/1.5 NA objective. Image acquisition and system control was done using MetaMorph Premiere software (Molecular Devices, LLC, San Jose, CA). Images were deconvolved with an iSIM-specific commercial plugin from Microvolution (Cupertino, CA) in FIJI. Live cell imaging microscopy experiments were performed on epifluorescent Leica DMi8 AFC microscope (Leica, Germany) equipped with a temperature (37°C), CO_2_ (5%), and humidity-controlled chamber (OkoLab, Italy) at 20✕, 40X and 63X magnifications. All analyses were performed automatically and/or manually utilizing Leica LAS X software (Leica, Germany) and the ImageJ/FIJI. Figures were composed using unmodified LAS X -generated TIFF images with Adobe Illustrator CC 2021 (Adobe).

### Automated detection and tracking of cells in collagen matrix

The detection of individual cell tracks and speeds in phase-contrast time-lapse images was performed with the following steps: i)correction of the background illumination, ii) detection of cell edges, iii) rescaling and applying masks, iv) watershed segmentation, v) tracking, and vi) filtering noise and correction for trajectory fragmentations.

The following steps allow us the detection of individual cell tracks and speeds in phase-contrast time-lapse images. First, the variation of the background illumination across each image was corrected by subtracting the image smoothed by the Gaussian filter (with a standard deviation of 10 pixels) from the original image. Second, the resulting background-corrected images were processed with the local standard deviation (STD) filter (with the 9x9-pixel neighborhood matrix) to highlight cell edge regions. Third, the resulting STD maps were smoothed by the Gaussian filter (with a standard deviation of 5) to obtain the gray-scale representation of cells. Next, the resulting images were re-scaled with a log function and masked with a threshold value of 6.4. To identify individual cells, the resulting binary masks were overlaid with the smoothed STD maps and subjected to our in-house watershed segmentation routine (using an algorithm that includes ‘draining’ criteria allowing to avoid over-segmentation (Hladyshau et al., 2021), which is typical for the traditional watershed approaches). Centroids of all 8-connected objects in the segmented images were recorded for tracking.

The tracking of cell centroids was performed based on minimal distance matching in consecutive time frames (within the radius of 40 pixels). The tracking pipeline includes the following steps. If two or more trajectories meet (e.g., one of the cells moves over or under another cell in the 3D matrix), the new position is added to the longest up to the time point track while the other one(s) ends. If a trajectory splits (e.g., another cell moves into the focus plane from above or below the tracked cell in the 3D matrix), the nearest new position is added to the current track, while the other new position starts a new track. Because cells move in the 3D environment, it is possible for cells to come in and out of the focus plane, which leads to recording several shorter tracks of the same cell. Since we do not use trajectory length as a phenotype characteristic, the occasional fragmentation of trajectories does not affect our analysis. However, it is also possible that a cell migrates at the border of the focus plane, repeatedly coming in and out of focus (i.e., flickering) and creating multiple ultra-short tracks. To reduce such disruptions of individual cell tracks, we apply a ‘flickering correction’ algorithm (Tsygankov et al., 2014) that detects co-localization of the end of one trajectory (*x_e_*,*y_e_*) at time *t* and the beginning of another trajectory (*x_b_*,*y_b_*) at time *t*+2 and merge these trajectories into one by assigning cell position at ((*x_e_*+*x_b_*)/2,(*y_e_*+*y_b_*)/2) at time *t*+1. Because we track cells with complex (ball-chain) shapes, it is possible for our detection algorithm to identify more than one compartment of the same cell as separate cells. As a result, a single cell may produce several nearly co-localized trajectories. To avoid a potential bias due to such double counting, we apply a ‘cloning correction’ algorithm that detects each trajectory remaining in the vicinity of another longer trajectory for the whole duration of this shorter track and removes such tracks from the record. Finally, we remove all the ‘tracks’ of immobilized debris in the image by disregarding trajectories that remain within the detection radius (40 pixels) for the entire track duration.

### Statistical analysis

Only pairwise comparisons as one-sided *t* tests between a control group and all other conditions are utilized to analyze the data, as well as between paired -Blebbistatin and +Blebbistatin co-treatment groups. Statistical analysis is performed using either KaleidaGraph 5 for MacOS (Synergy Software) or Prism 7b (GraphPad Software, Inc). The exact p values are indicated on the plots, unless the p<0.0001, *i.e.* below the cut-off lower limit for Kaleidagraph and Prism software. Sample size n for each comparison is reported in the corresponding plots (*i.e.* n reflects the number of measured individual cells). Number of replicates is 3, unless specified otherwise. Data are shown as box and whiskers diagrams: first quartile, median, third quartile, and 95% percent confidence interval.

## DATA AVAILABILITY STATEMENT

The authors declare that the data supporting the findings of this study are available within the paper, the supplementary information, and any data can be made further available upon reasonable request.

## ACKNOWLEDGEMENTS

E.D.T. and this work were supported by the Department of Pharmacology, Penn State College of Medicine *via* the startup funds. D.T. is supported by the National Science Foundation (CMMI 1942561) and the National Institutes of Health (R01GM136892). A.S.Z., X.M., and A.M. were supported by the FDA Intramural Research Program of the Center for Biologics Evaluation and Research. A.X.C.R. and C.S. were supported by the National Institutes of Health (NIH) Intramural Research Program in the National Institute of Biomedical Imaging and Bioengineering (NIH grant # ZIA EB000094) and by the NIH Distinguished Scholars Program. P.J.S received the support of Human Frontier Science Program (HFSP) RGP 0032-2022 and Forschungszentrum Medizintechnik Hamburg (FMTHH, grant 04fmthh2021). We thank Christian Combs and Daniela Malide for the Light Microscopy Core support at the National Heart, Lung, and Blood Institute, NIH.

## MOVIES LEGENDS

**Movie 1**. Brightfield live video-microscopy sequence of an hCD4+ T cell migrating through 3D collagen type I matrix.

**Movie 2**. Structured illumination microscopy. Fluorescent visualization of the hCD4+ T cell’s nucleus passage (Hoechst, cyan) through the cytoskeletal ring (SiR-actin, yellow) that constitutes a key peristaltic translocation event during T cell’s amoeboid locomotion within 3D collagen matrix (magenta).

**Movie 3**. Epifluorescent microscopy. F-actin-dense rings (side views, seen as bands, SiR-actin live staining, yellow) form a cortical tube that peristaltically channels the nucleus (blue) *via* amoeboid passage as seen in hCD4+ T cells.

Supplementary Information (Key resources table):

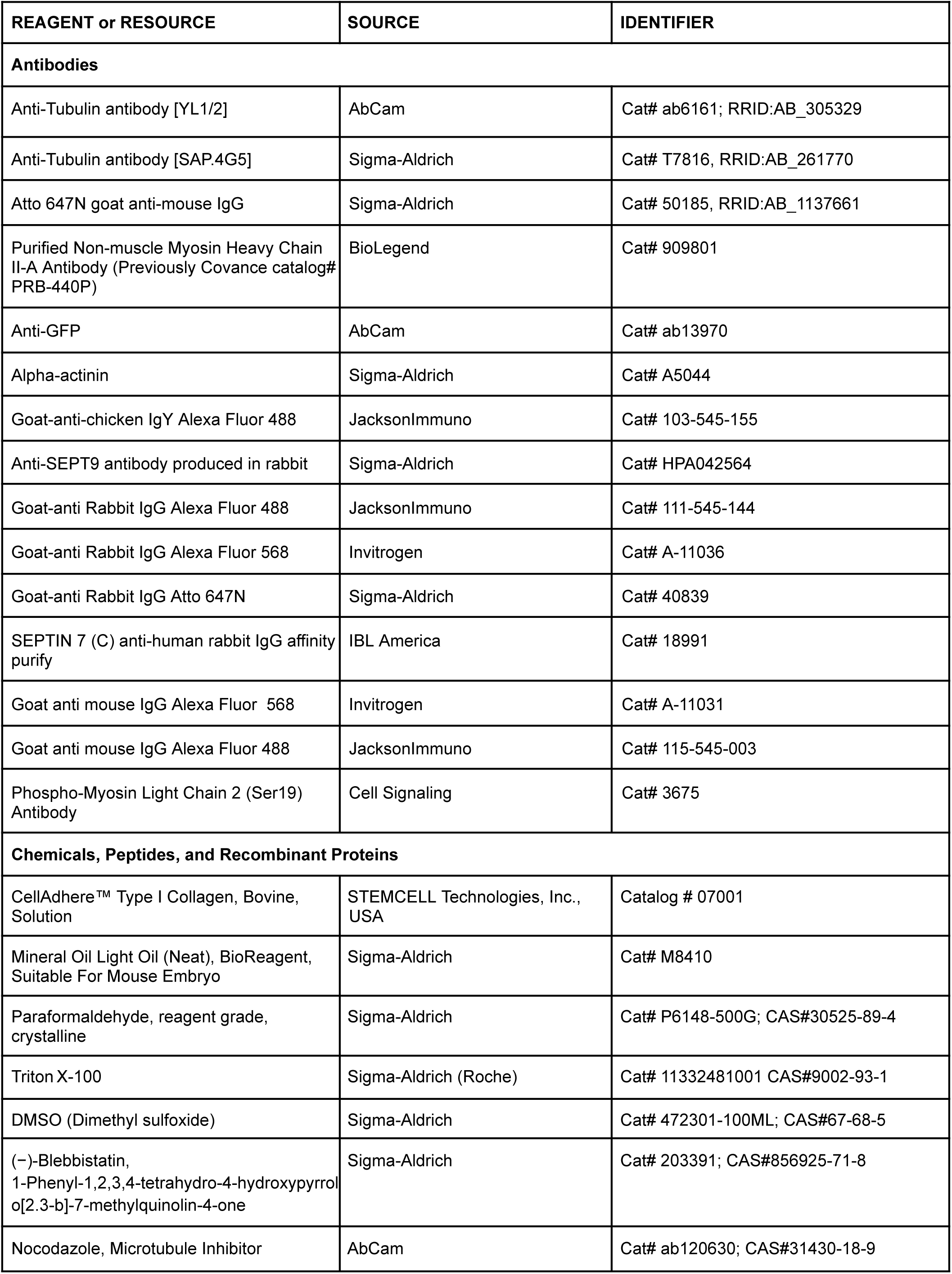
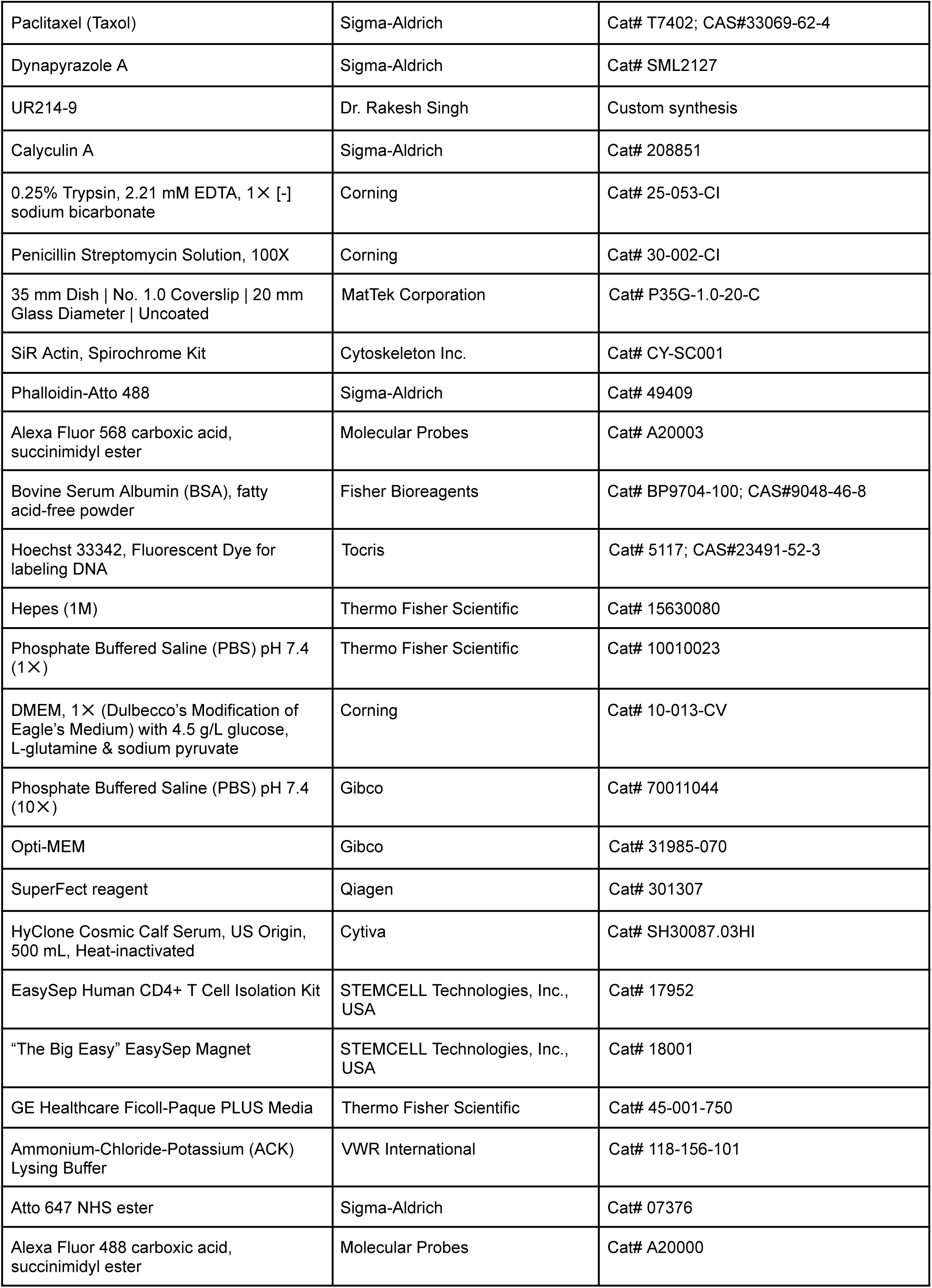
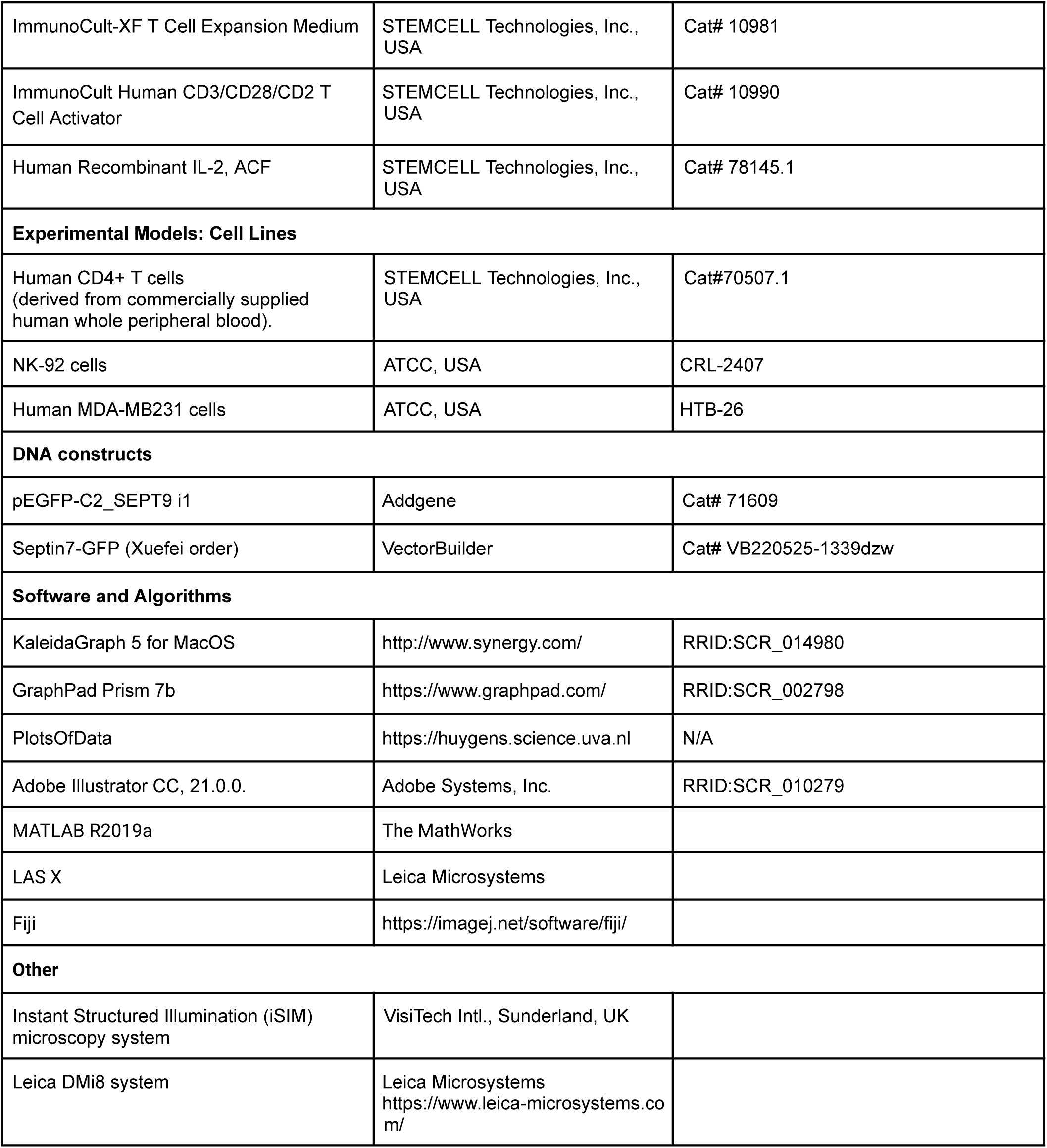

---

**Supplemental Figure SI1.**
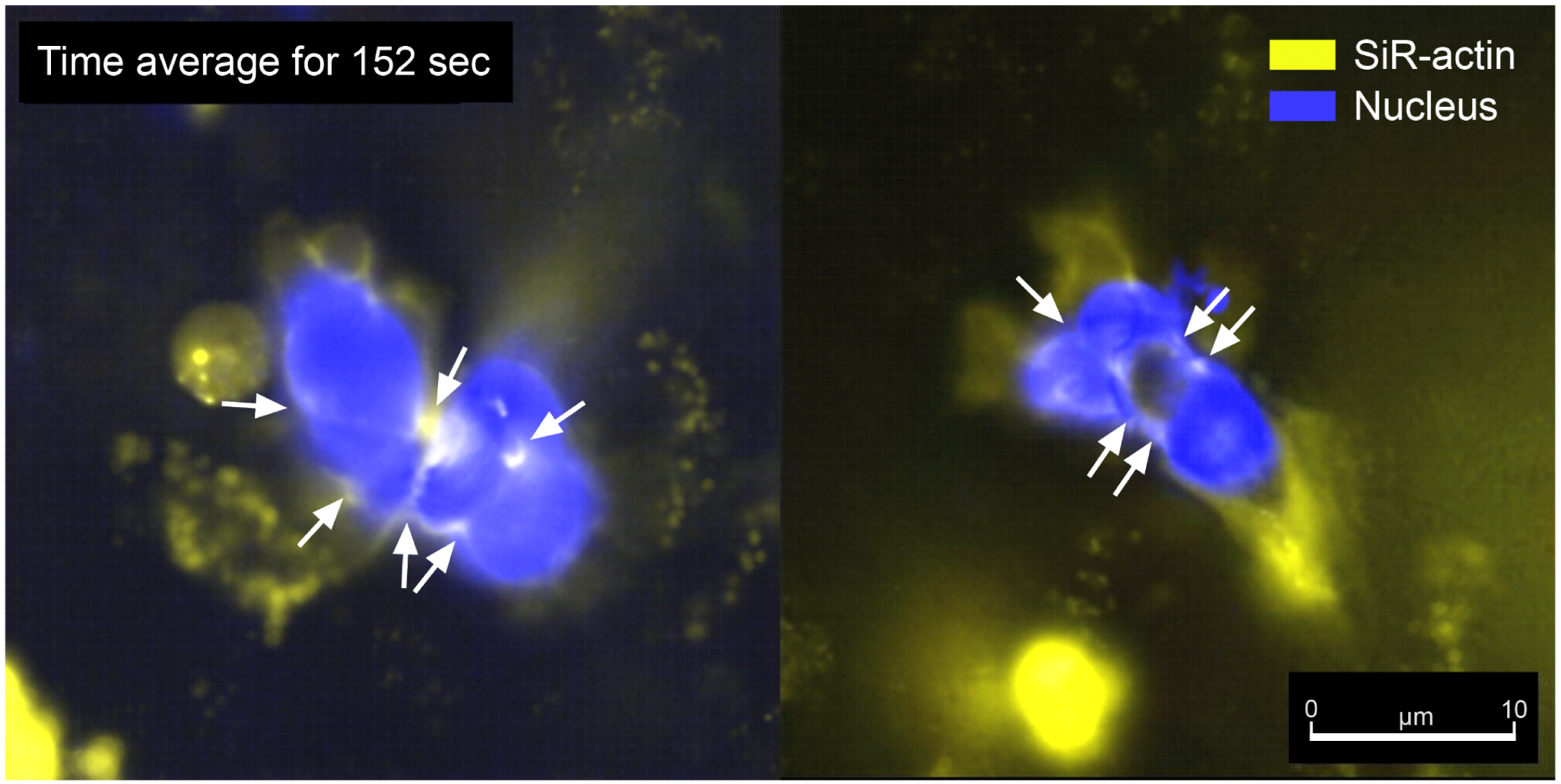
Time average projections of the migrating hCD4+ T cells within the collagen matrix reveal stationary cortical rings and the peristaltic translocation of the nucleus. F-actin-dense rings (*arrows*, side views, seen as bands, SiR-actin live staining) form a cortical tube that channels the nucleus (blue) *via* peristaltically amoeboid cortex passage. *The shown time average projections correspond to the video sequences in Movie 3*.

**Supplemental Figure SI2.**
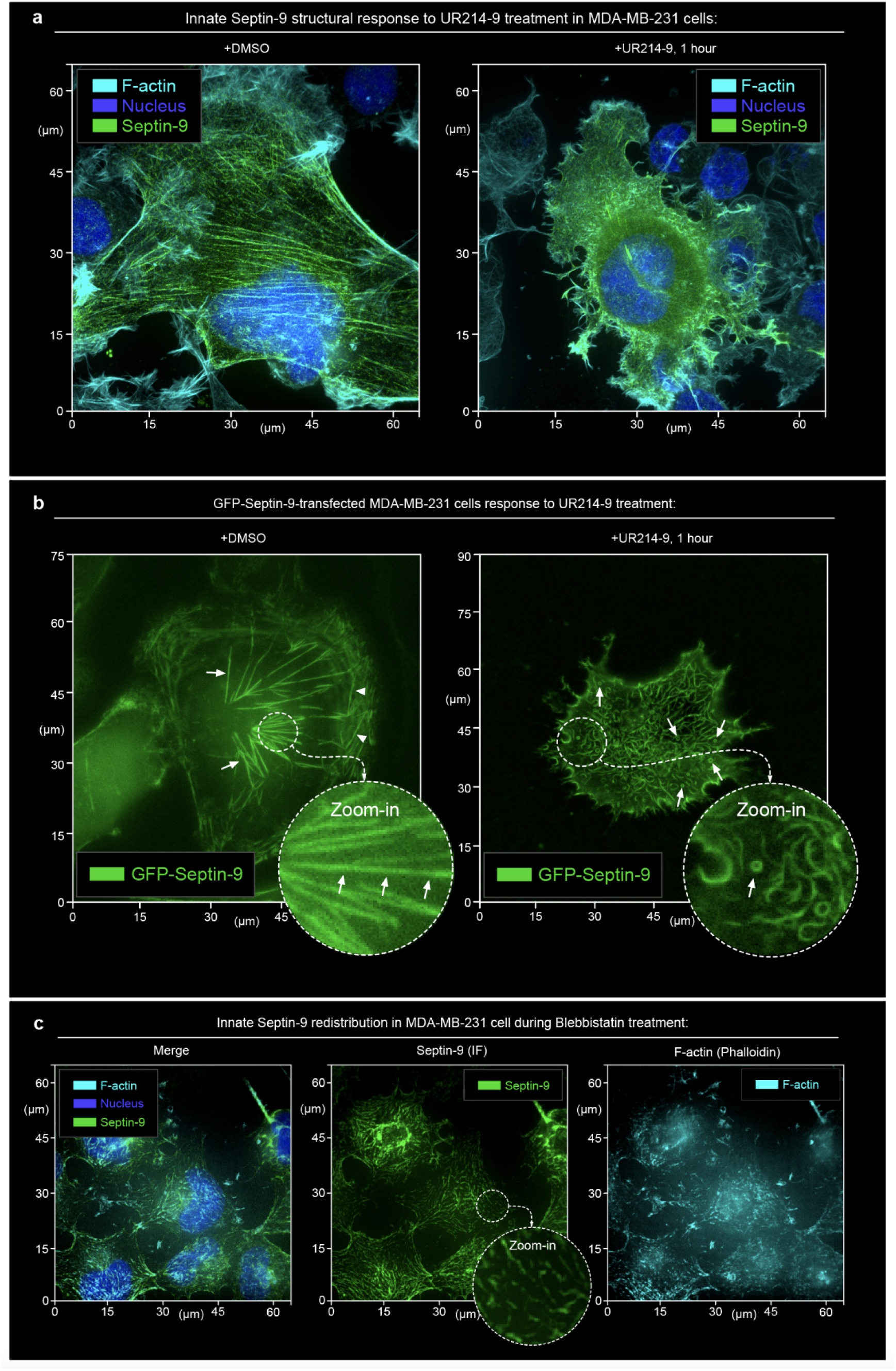
Septins (Septin-9 staining and live imaging) colocalize with tensile actomyosin structures (stress-fibers) that disintegrate both in the presence of septin inhibitor (+UR214-9) and during suppression of actomyosin contractility (+Blebbistatin). **(a)** Septin filaments disassemble upon the UR214-9 treatment in MDA-MB-231 cells, as shown by Septin-9 staining. MDA-MB-231 cells, spread on a glass surface form stress-fibers (SiR-Actin) and septin filaments (Septin-9 immunofluorescence) as a single cytoskeletal network **(+DMSO)**. Cell treatment with a septins inhibitor results in the dissociation of the actin and septin filaments and retraction of cells **(+UR214-9,** 1 hour). Instant structured illumination microscopy. **(b)** *Left* - GFP-Septin-9 overexpressing and fully spread MDA-MB-231 cells show Septin-9 filaments in radial dorsoventral stress-fibers and within the transverse arcs **(+DMSO)**. *Right* - septin inhibitor induces septin filaments disintegration **(+UR214-9, 1 hour)**. **(c)** Septins (Septin-9) and F-actin (phalloidin) filaments disintegrate upon inhibition of actomyosin contractility **(+Blebbistatin)**.

**Supplemental Figure SI3.**
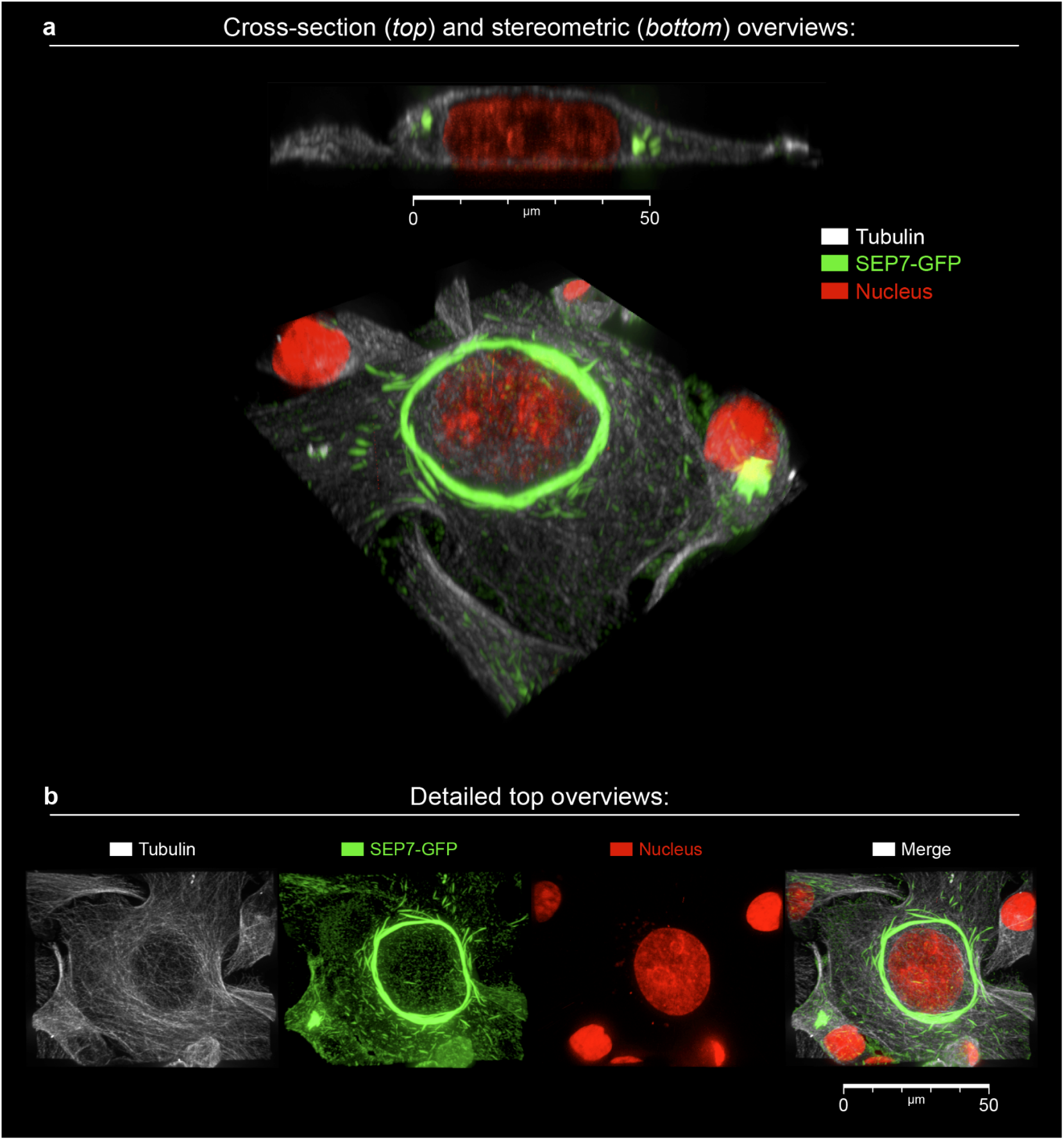
Cell surface curvature-sensitive assembly of septin-7-GFP) filaments, conforming along the inner side of cortex at the perinuclear convex ridge in the MDA-MB-231 cells. **(a)** Cross-section (*top*) and stereometric (*bottom*) views of the MDA-MB-231 cell, transiently transfected with human SEP7-GFP, spread on the glass surface. **(b)** Detailed top overviews of key components of MDA-MB-231 upon formation of SEP7-GFP rings.

